# Dose and sex-specificity in Canagliflozin-mediated neuroprotection in aging mice

**DOI:** 10.64898/2026.07.02.734998

**Authors:** Dulmalika Herath Manchanayake, Hashan Jayarathne, Sydney Scofield, Naduni Dasanthi Hitihami Mudiyanselage, Lindsey DeHaan, Omar Kadri, Nabeelah Rouf, Brett C. Ginsburg, Richard A. Miller, Marianna Sadagurski

## Abstract

Canagliflozin (Cana), an SGLT2 inhibitor prescribed for type 2 diabetes, extends median lifespan by 14% in male but not female UM-HET3 mice at 180 ppm, with male-specific neuroprotective effects, despite females accumulating higher drug concentrations in blood and brain. Here, we tested whether reducing the dose to a subclinical level of 60 ppm could provide neuroprotective benefits in females by reducing drug accumulation. Starting treatment at 7 months of age, Cana at 60 ppm improved glucose tolerance in both sexes at 18 months and increased water and food intake, consistent with SGLT2 inhibition, but produced only a transient reduction in fat mass in males after one month on diet, with no sustained effect on body weight in either sex. At 60 ppm, Cana did not improve cognitive function at 18 months or reduce neuroinflammation in males, whereas females showed reduced hippocampal microgliosis and astrogliosis at 24 months. Pharmacokinetic analysis demonstrated that females accumulated 3-to 5-fold higher Cana concentrations than males across brain regions, blood, and liver. Together, these findings demonstrate that neither dose reduction nor greater drug accumulation drives neuroprotective benefit in females, indicating fundamental sex differences in the biological response to SGLT2 inhibition and suggesting that the sex-specific longevity effects of Cana are not simply a matter of dose.

## INTRODUCTION

Aging is the primary risk factor for cognitive decline and neurodegenerative disease, with sex differences shaping the progression of brain aging. At the same time, the sex-specific responses to pharmacological intervention remain incompletely understood and understudied. Among the most promising anti-aging interventions to emerge from preclinical testing is Canagliflozin (Cana), a sodium-glucose co-transporter 2 inhibitor (SGLT2i) originally developed for the treatment of type 2 diabetes (Vallon 2015). The NIA Interventions Testing Program (ITP) demonstrated that Cana, administered at 180 ppm in the diet, extends median lifespan by 14% in genetically heterogeneous UM-HET3 male mice, with no significant lifespan benefits in females, while improving glycemia and body weight in both sexes (Miller *et al*. 2020). In addition to Cana, the ITP has identified several interventions with sex-specific effects, including Acarbose, which preferentially extends lifespan in males, and 17-α-Estradiol, which benefits males but not females (Harrison *et al*. 2014; Garratt *et al*. 2018); however, unlike other interventions, Cana is widely used in clinics, making the question of why it benefits males but not females more clinically relevant for aging.

We previously demonstrated that chronic Cana treatment at 180 ppm significantly enhanced central insulin sensitivity in the hypothalamus and hippocampus of aged male mice, reduced age-associated neuroinflammation, and improved locomotor activity in aged male, but not female mice (Jayarathne *et al*. 2022). Pharmacokinetic analysis revealed that females accumulate significantly higher concentrations of Cana in the brain and blood than males, indicating that the sex difference in neuroprotective efficacy is not due to insufficient drug exposure in females. Comparison of cortical Cana concentrations between doses confirmed dose-dependent accumulation in both sexes, with higher concentrations at 180 ppm, consistent with the dietary dose difference. (Miller *et al*. 2020). Subsequent transcriptomic, proteomic, and metabolomic profiling of the aging hippocampus revealed that Cana induced mitochondrial function, insulin signaling, and cyclic GMP-dependent signaling in males, with improvements in hippocampal-dependent learning and memory in aged males at 14 months, but not females. At the same time, in females, benefits were limited across all molecular layers (Jayarathne *et al*. 2025). Similarly, at the hypothalamic level, Cana treatment increased energy expenditure, with sex-specific transcriptional remodeling of hypothalamic metabolic circuits in aged males but not females (Jayarathne *et al*. 2024). Together, these findings demonstrate that the sex-specific effects of Cana on brain aging are not simply explained by differential drug accumulation, pointing to intrinsic differences in CNS responsiveness between males and females.

A recent study found that initiating Cana at 180 ppm at 16 months of age extended male lifespan but shortened female lifespan by 6%. Importantly, blood levels of Cana were significantly higher in aged females than in young males, indicating sex-specific differences in drug accumulation as a potential mechanism for this female-specific response (Miller *et al*. 2024). Taken together, these data suggest that Cana at 180 ppm is beneficial to males and neutral to females when started early, with both sexes showing metabolic improvements, but detrimental to females when started late. This sex-specific response raises two mechanistic possibilities: either 180 ppm produces off-target or counterproductive effects in female tissues, that are amplified at higher drug exposures, or females require a different dose to drive the protective pathways that benefit males without the adverse consequences of drug accumulation. Recently, the ITP initiated a new study to test the effects of Cana at 60 ppm (one-third of the original dose) on lifespan, starting at 7 months of age. Here, we characterize the metabolic, pharmacokinetic, and neuroprotective effects of this lower dose in both sexes and specifically test whether reducing the dose can provide beneficial CNS effects in female mice.

## MATERIALS AND METHODS

### Animals

Genetically heterogeneous UM-HET3 mice, produced by the four-way cross [(BALB/cByJ x C57BL/6J)F1 x (C3H/HeJ x DBA/2J)F1], were obtained from the University of Michigan ITP colony. Mice were housed under standard conditions (12h light/dark cycle, 22°C, ad libitum food and water) in the Wayne State University animal facility. All procedures were approved by the Wayne State University Institutional Animal Care and Use Committee. Mice are housed at three males or four females per cage from weaning and are provided food (TestDiet 5LG6: 17.5% protein, 5.6% fat) and water ad libitum. Mice in the Cana group (5LG6 w/10% 60 ppm Cana) received this agent at 60 mg per kg of chow, from 7 months of age, and sacrificed at 24 months of age as previously described (Miller *et al*. 2020). Body weight was monitored monthly. Body composition (fat mass and lean mass) was measured by Echo-MRI.

### Glucose tolerance testing

Intraperitoneal glucose tolerance testing (IPGTT) was performed at 12 and 18 months of age. Mice were fasted for 6 hours before testing. Glucose (2 g/kg body weight) was administered by intraperitoneal injection. Blood glucose was measured from tail vein blood at 0, 15, 30, 60, 90, and 120 minutes post-injection using a glucometer (Contour Next, Bayer).

### Indirect calorimetry

Metabolic parameters were assessed at 24 months of age using the PhenoMaster automated metabolic cage system (TSE Systems). Mice were individually housed and allowed to acclimate for 48 hours before a 72-hour recording period. Oxygen consumption (VO_2_), carbon dioxide production (VCO_2_), respiratory exchange ratio (RER), energy expenditure (EE), food intake, and water intake were continuously recorded. Data were analyzed by CalR (https://calrapp.org/) (Mina *et al*. 2018).

### Perfusion and immunolabeling

Mice were anesthetized and perfused using phosphate buffer saline (PBS) (pH 7.5), followed by 4% paraformaldehyde. Brains were postfixed, dehydrated, and then sectioned coronally (30□μm) using a sliding microtome, followed by immunofluorescent analysis as described (de Lima *et al*. 2021). For immunohistochemistry brain sections were washed with PBS six times, blocked with 0.3% Triton X-100 and 3% normal donkey serum in PBS for 2 h; then the staining was carried out with the following primary antibodies overnight: rabbit anti-GFAP (1:1000; Millipore, Cat. No. ab5804) and goat anti-Iba1 (1:1000 Abcam Cat. No. ab5076). After the primary antibody, brain sections were incubated with AlexaFluor-conjugated secondary antibodies for 2 h (Invitrogen). Microscope images of the stained sections were obtained using a Nikon 800 fluorescent microscope with Nikon imaging DS-R12 color-cooled SCMOS, version 5.00.

### Quantification

For cell quantification and immunoreactivity analysis, images were acquired from at least 3 sections containing the hypothalamus and hippocampus from each brain, spanning bregma −0.82 to −2.4□mm (according to the Franklin mouse brain atlas). Serial brain sections were made at 30□μm thickness. All sections were arranged from rostral to caudal to assess the distribution of labeled cells from the equivalent sections for all stains. Fiji-ImageJ was used to count GFAP- and Iba1-positive cells and to measure immunoreactivity. All microscopy images and quantifications were performed by investigators who were blinded to the sample’s ID.

### Behavioral Assays

All behavioral testing was conducted with a 2-day rest period between tests. The tests were conducted in the following order: open field test, spontaneous alteration, rotarod, grip strength, accelerated rotarod, novel object recognition test, and Barnes maze. On each test day, animals were transported from the housing room to the procedure room and left to acclimate for at least 1□h. Animals were provided with ad libitum water and food in the procedure and housing rooms. Silence in the room was maintained throughout testing to avoid any disturbance or stress that might affect the animal’s natural behavior. Between tests, the arena was sanitized with 70% ethanol and dried before introducing the next animal. Lighting in the testing room was consistent with the housing room, except for the novel object test, which was conducted under ambient lighting (∼20–35□lx). Animals’ movement within the arenas was recorded using a 22 Series CMOS camera, and the recorded videos were analyzed using the Any-Maze video tracking system V 7.49. A brief description of each behavioral test is provided below.

### Open Field Test

The open field arena was 31.7″□×□31.7″□×□11.6″ and made with polyvinyl chloride (PVC). The center of the arena was defined as a 16″□×□16″ section in the center, and the four corner zones were also tracked. The animals inside the arena were recorded for 10□min. As the test started, animals were placed in the center of the arena and allowed to walk freely. After each test, the animals were returned to their home cage. The travel distance and time spent in the center for each animal were calculated using the Any-maze software.

### Spontaneous Alternation Test (Y Maze)

The Y-maze arena was 13.5″□×□2″□×□7.8″ and made with polyvinyl chloride (PVC). The three arms were labeled as “A,” “B,” and “C” for later analysis purposes. Animals were placed halfway into the start arm (A), facing the center of the Y, and allowed to explore the Y-maze for 8□min. The sequence of entries into each arm was recorded. Using the Any-maze video tracking software, the % of spontaneous alterations was calculated as before (Jayarathne *et al*. 2025).

### Barnes Maze

The Barnes Maze arena was 18″ in radius with 20 holes around the arena’s edge and four distinct visual cues. The test was conducted in three phases: the acquisition phase, the short-term memory (STM) test, and the long-term memory (LTM) test. On the first day before the acquisition started, animals were introduced to the arena, guided to the escape hole, and allowed to remain inside for 2□min. Then, three acquisition trials were conducted for each day of the acquisition phase, with a maximum time of 180□s for each trial. During the acquisition phase, the location of the escape hole and visual cues were not changed. On the 5th day, the STM test was performed, and on the 12th day, the LTM test was performed. During the STM and LTM tests, the escape hole was removed, and the animal was introduced to the middle of the arena and allowed 90 s to locate the escape hole. All these phases were conducted while white noise was in the background. Latency to the target hole and time spent at the target hole were analyzed using Any-maze video-tracking software.

### Pharmacokinetic analysis

Tissue Cana concentrations were measured by LC-MS/MS. At 24 months, mice were euthanized, and the hypothalamus, hippocampus, cortex, and liver were rapidly dissected, weighed, snap-frozen in liquid nitrogen, and stored at -80°C. Tissue was homogenized in ice-cold extraction buffer, and Cana was extracted by protein precipitation with acetonitrile containing an internal standard. Samples were analyzed on a triple quadrupole mass spectrometer (Waters TQ-S) coupled to a UPLC system. Cana concentrations were determined from standard curves and normalized to tissue weight.

### Statistical analysis

Data sets were analyzed using Student’s t-test and two-factor analysis of variance (two-way ANOVA) with the general linear model function and a full factorial model that included the effects of treatment, sex, and the interaction between sex and treatment, followed by a Newman–Keuls post hoc test. All data were presented as mean□±□SEM. p <□0.05 was considered significant. TIBCO Statistica® v. 13.5.0.17 was used for statistical analysis.

## RESULTS

### Low-dose Canagliflozin improves glucose tolerance in male and female UM-HET3 mice

UM-HET3 males and females were placed on a 60 ppm Cana diet beginning at 7 months of age and were assessed longitudinally through 24 months. One month after diet initiation, Cana-fed males showed a significant reduction in fat mass compared with controls, with no significant change in lean mass (Fig. 1A-C). Female mice, however, showed no significant changes in body weight, fat mass, or lean mass after one month on Cana (Fig. 1A-C). Longitudinal assessment of body weight from 7 to 22 months of age revealed that both Cana-fed male and female mice maintained slightly, but not significantly, lower body weights than the control groups at all time points (Fig. 1D and E). Glucose tolerance at 12 months of age showed no significant differences in 60 ppm Cana-fed mice in either sex (Fig. 1F and G). However, at 18 months, 60 ppm Cana significantly improved glucose tolerance in both male and female mice (Fig. 1H and I), suggesting that Cana has beneficial effects on peripheral tissue metabolism at low doses in both sexes during aging.

**FIGURE 1:**
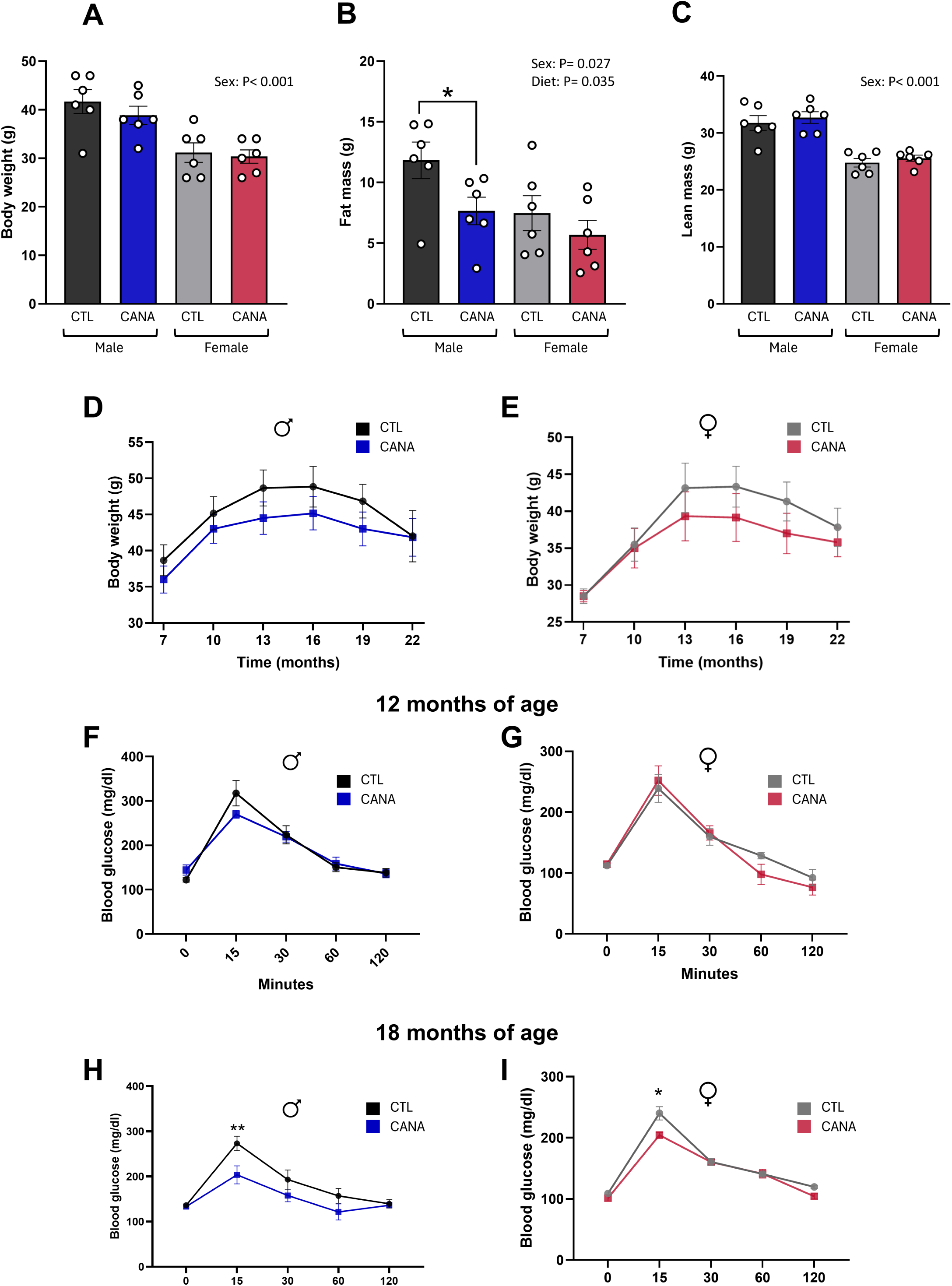
Body composition and glucose homeostasis in UM-HET3 mice fed 60 ppm Canagliflozin. (A–C) Body weight (A), fat mass (B), and lean mass (C) in male and female mice at 1 month after diet initiation. (D, E) Longitudinal body weight from 7 to 22 months of age in males (D) and females (E). (F, G) Blood glucose during the intraperitoneal glucose tolerance test (IPGTT) at 12 months of age in males (F) and females (G). (H, I) Blood glucose during IPGTT at 18 months of age in males (H) and females (I). Data are presented as mean ± SEM. *p < 0.05, **p < 0.01. Two-way ANOVA with Newman-Keuls post hoc test; p-values for main effects of sex and diet are indicated.

### Low-dose Canagliflozin increases water and food intake in aged mice without changes in energy expenditure

At 24 months of age, Cana-fed mice of both sexes exhibited significantly increased water intake during the dark cycle and increased food intake over the 24-hour period compared with controls (Fig. 2A–D, M, N), consistent with the glucosuria effects of SGLT2 inhibition, which promote compensatory increases in fluid and caloric intake (Tang *et al*. 2022). However, no significant differences in energy expenditure (EE) or respiratory exchange ratio (RER) were observed in either sex (Fig. 2E-L).

**FIGURE 2:**
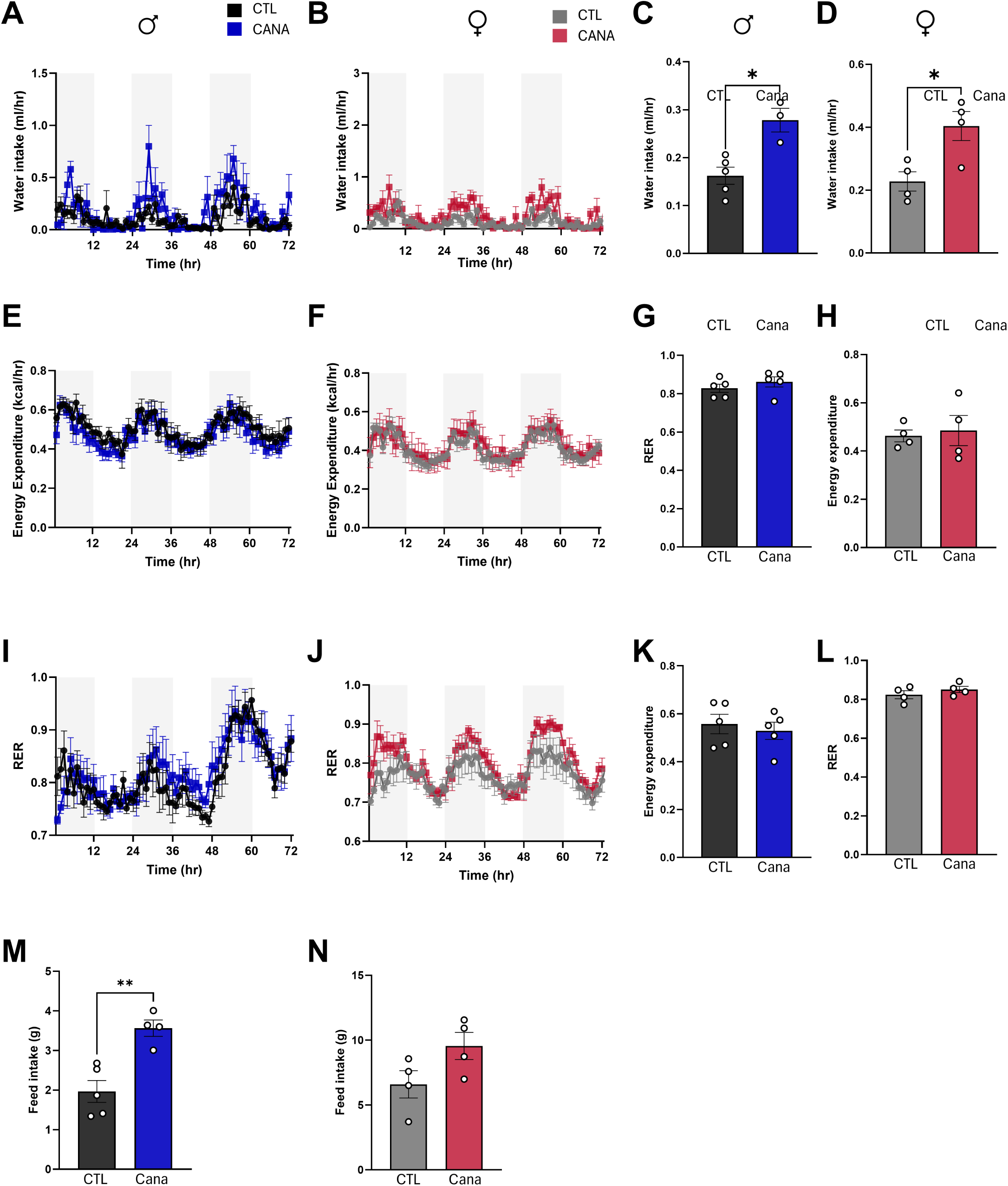
Metabolic parameters in 60 ppm Cana-fed UM-HET3 mice at 24 months of age. (A, B) Water intake over 72 hours in males (A) and females (B), average dark cycle water intake (C, D). (E, F) Energy expenditure (EE) in males (E) and females (F), average dark cycle EE (G, H). (I, J) Respiratory exchange ratio (RER) in males (I) and females (J), average dark cycle RER (K, L). (M, L) Food intake over 24h (for 3 days) in males (M) and females (N). Data are presented as mean ± SEM. *p < 0.05. Two-way ANOVA with Newman-Keuls post hoc test; p-values for the main effect of diet during the dark cycle are indicated.

### Sex-specific accumulation of Canagliflozin in brain regions and liver

Cana contents in blood plasma, hypothalamus, hippocampus, cortex, and liver were measured by high-performance liquid chromatography tandem mass spectrometry as before (Miller *et al*. 2020; Jayarathne *et al*. 2025). Cana was detectable in all brain regions, liver, and blood in both sexes, confirming its penetration into the brain and peripheral organs even at the lower dose. Although plasma Cana concentrations tended to be higher in females than in males, this did not reach statistical significance (Fig. 3A). Female mice showed significantly higher Cana concentrations in the hippocampus, cortex, and liver compared to males (Fig. 3D, 3E, 3B). In contrast, hypothalamic Cana concentrations did not differ significantly between sexes (Fig. 3C). Comparison of Cana cortical concentrations between the 60 ppm and 180 ppm doses revealed dose-dependent differences in both sexes, with higher cortical concentrations at 180 ppm (Fig. 3 F).

**FIGURE 3:**
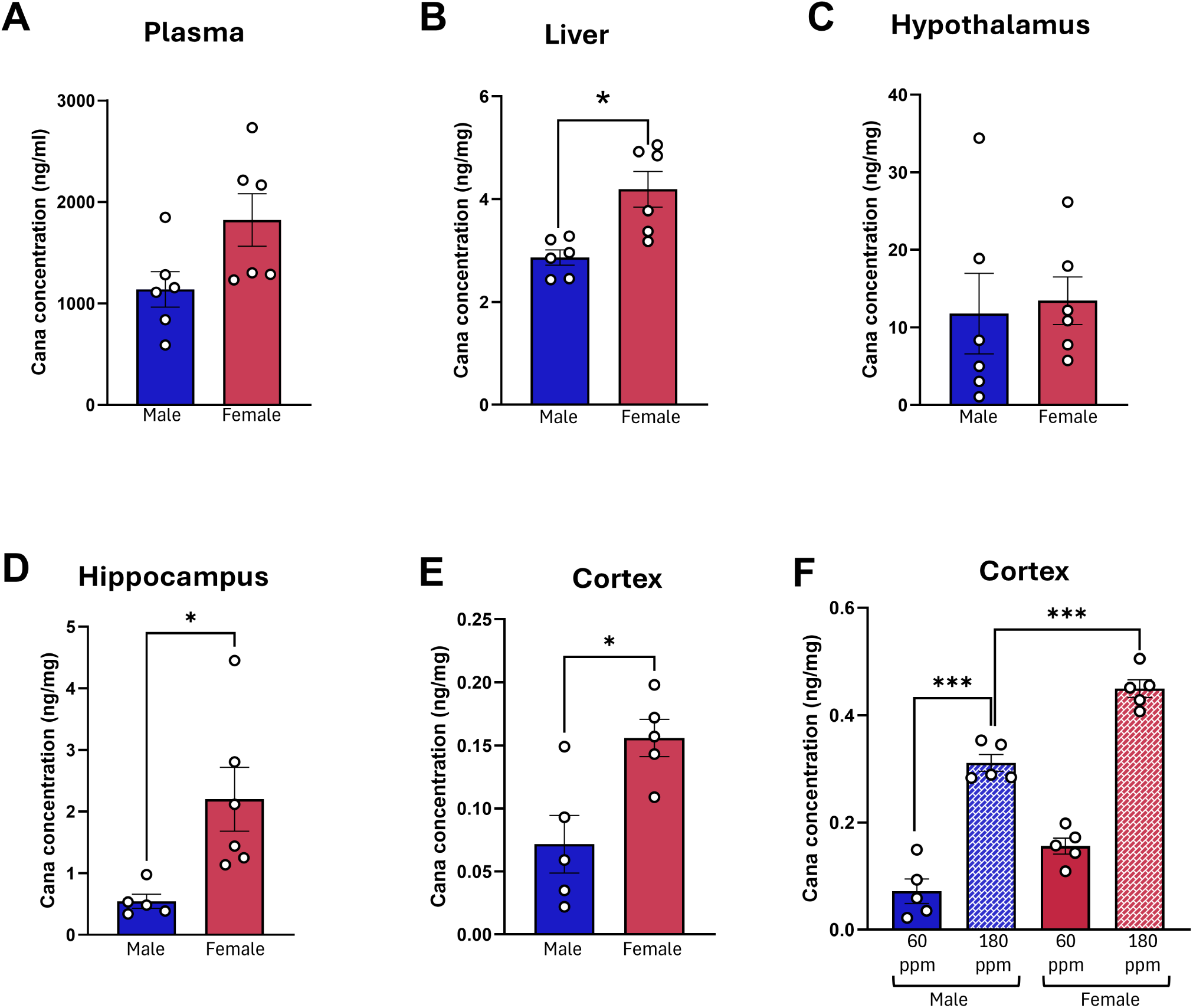
Sex-specific accumulation of Canagliflozin in brain regions and liver. Cana concentrations measured by LC-MS/MS in plasma (A), liver (B), hypothalamus (C), hippocampus (D), cortex (E) of male and female mice fed a 60 ppm Cana diet and cortex (F) of male and female mice fed 180 or 60 ppm Cana diet. p values were determined by unpaired Student’s t-test (A–E) or two-way ANOVA with Newman-Keuls post hoc test (F). *p < 0.05, **p < 0.01, ***p < 0.001.

### Low-dose Canagliflozin does not improve motor or cognitive function in aging UM-HET3 mice

A comprehensive battery of behavioral assays was conducted at 18 months of age. Rotarod performance (Fig. 4A) and grip strength (Fig. 4B) were not significantly different between Cana and control groups in either sex. Likewise, the novel object recognition discrimination index (Fig. 4C), spatial working memory, assessed by spontaneous alternation in the Y-maze (Fig. 4D), and the open field test (Fig. 4E) did not differ significantly between the Cana and control groups in either sex. In the Barnes maze, male Cana-fed mice showed a trend toward reduced escape latency during the acquisition phase. Still, neither short-term nor long-term memory probe trials showed significant differences between Cana-fed and control males (Fig. 4F-H). Female mice showed no differences in any Barnes maze parameter (Fig. 4F-H). These results contrast with the significant improvements in Barnes maze performance observed in males at 14 months with 180 ppm Cana (Jayarathne *et al*. 2025), demonstrating that the cognitive benefits of Cana are dose-dependent and absent in females at either dose.

**FIGURE 4:**
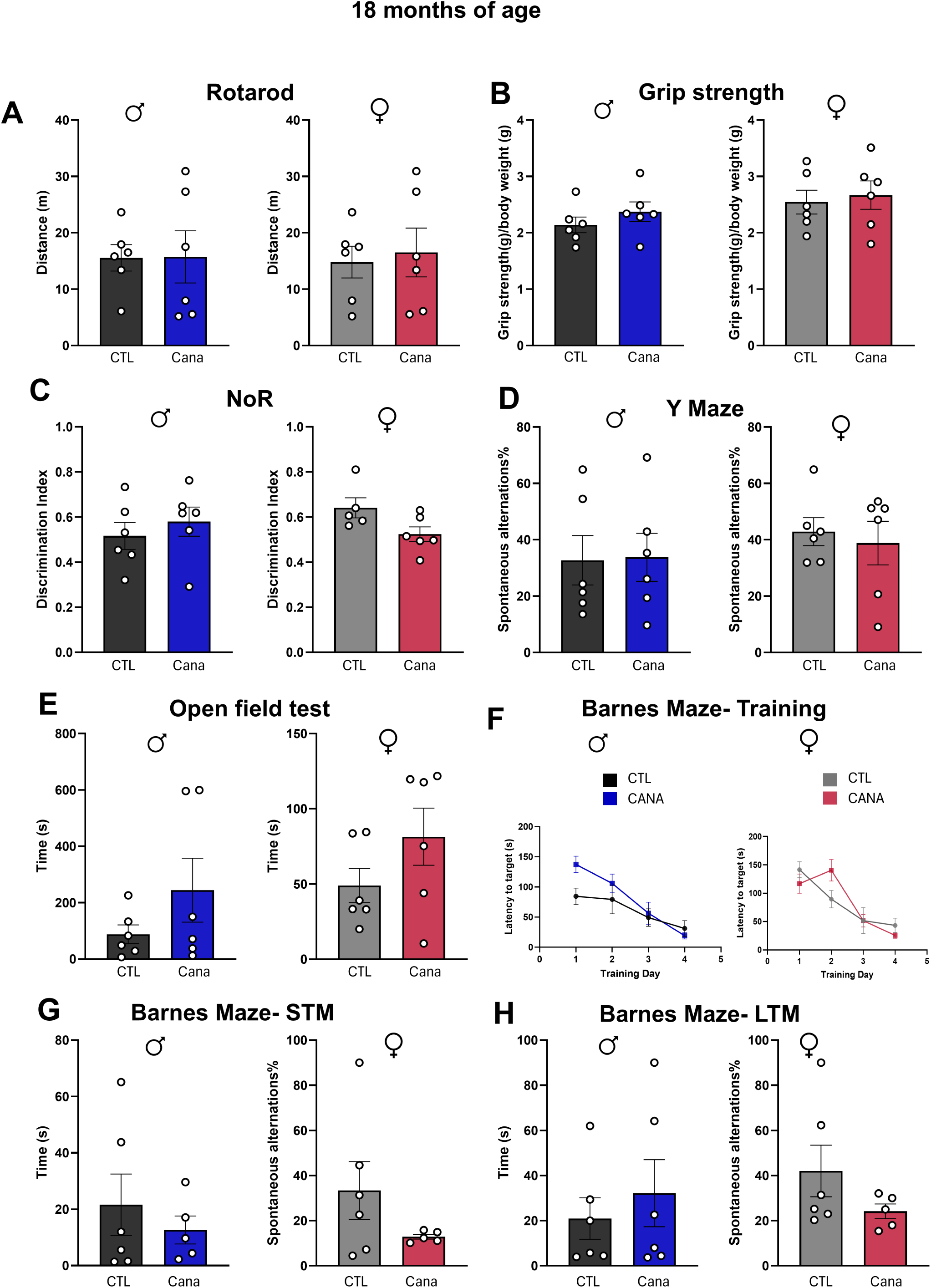
Cognitive and motor behavioral assessment in 60 ppm Cana-fed UM-HET3 mice at 18 months of age. (A) Rotarod performance was assessed as the distance traveled by males and females. (B) Grip strength in males and females. (C) Novel object recognition (NoR) discrimination index in males and females. (D) Spontaneous alternation percentage in the Y-maze in males and females. (E) Open field test distance traveled in males and females. (F) Barnes maze acquisition phase: latency to target hole across 5 training days in males and females. (G) Barnes maze short-term memory (STM) probe trial latency and (H) long-term memory (LTM) probe trial latency in males and females. Data are presented as mean ± SEM. Two-way ANOVA with Newman-Keuls post hoc test.

### Low-dose Canagliflozin produces region- and sex-specific effects on hippocampal neuroinflammation

Neuroinflammation in the hippocampus and hypothalamus was assessed at 24 months of age using immunofluorescence for GFAP, a marker of astrogliosis, and IBA1, a marker of microgliosis. At this age, both GFAP and IBA1 immunoreactivity are significantly elevated compared to young mice in UM-HET3 mice (Sadagurski *et al*. 2017). IBA1-positive and GFAP-positive cells in the arcuate nucleus of the hypothalamus (ARC) did not differ significantly between the treatment groups in either sex (Fig. 5A, B, G, and H). In the hippocampus, however, Cana-fed female mice showed significant reductions in IBA1-positive cells in both the CA3 subregion and the dentate gyrus (DG) compared to controls, indicating reduced age-associated microgliosis across hippocampal subregions (Fig. 5C-F). In addition, female mice showed a significant reduction in GFAP-positive cells in the DG, suggesting that 60 ppm Cana produces a female-specific reduction in hippocampal astrogliosis in this subregion (Fig. 5K, L). No significant differences in hippocampal IBA1 or GFAP immunoreactivity were detected in males at any hippocampal region examined (Fig. 5C-F and I-L). Notably, this female-specific hippocampal response was not observed at 180 ppm, where hippocampal anti-neuroinflammatory benefits were male-specific (Jayarathne *et al*. 2022), suggesting a dose-dependent response in the female hippocampus, in which low-level SGLT2 inhibition is beneficial while higher concentrations are not.

**FIGURE 5:**
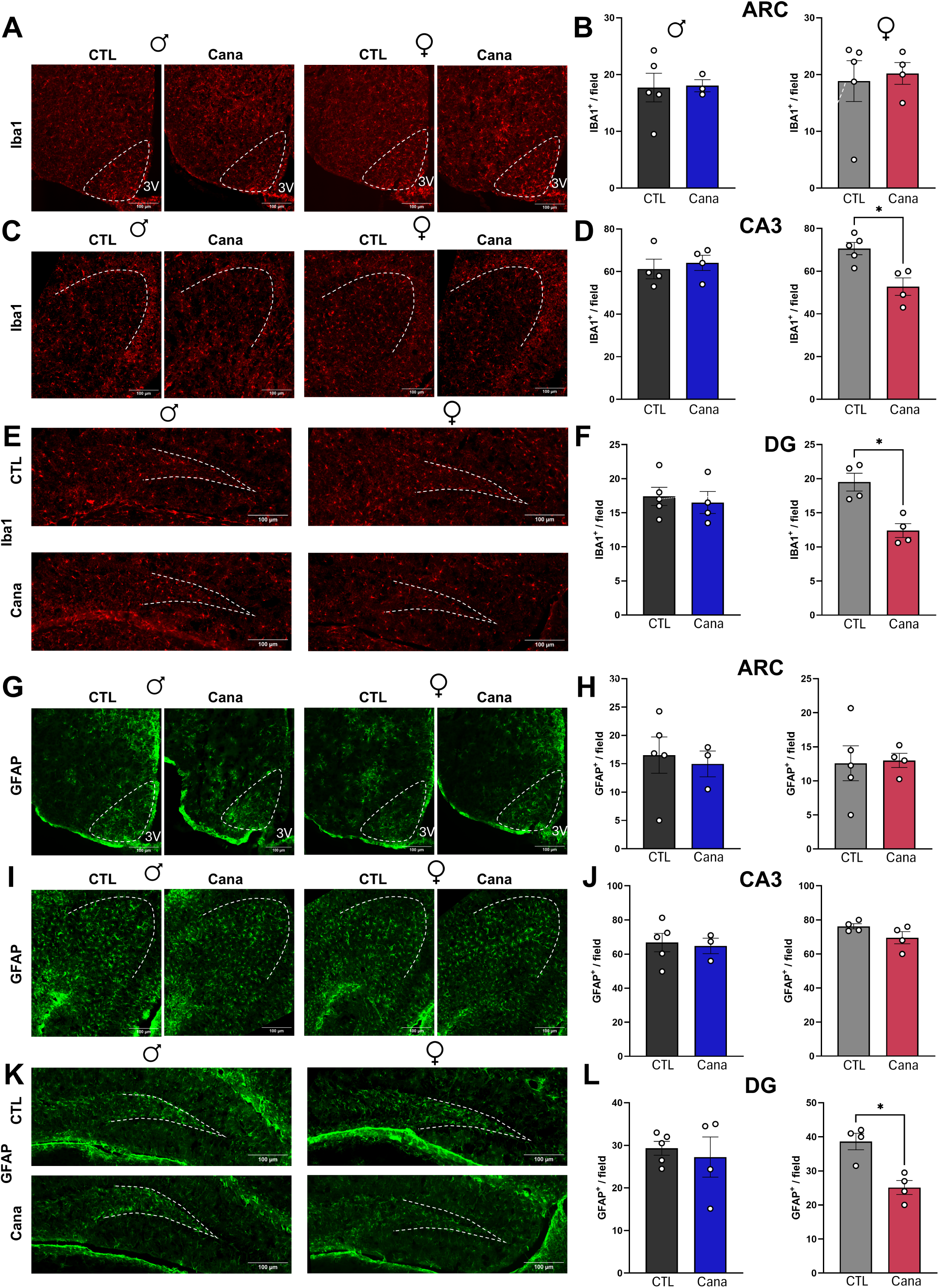
Region- and sex-specific effects of 60 ppm Canagliflozin on hippocampal and hypothalamic neuroinflammation at 24 months of age. (A–F) IBA1 immunofluorescence images and quantification in the arcuate nucleus (ARC) of the hypothalamus (A, B), CA3 subregion of the hippocampus (C, D), and dentate gyrus (DG) (E, F) in control and Cana-fed male and female mice. (G–L) GFAP immunofluorescence images and quantification in the ARC (G, H), CA3 (I, J), and DG (K, L). IBA1-positive and GFAP-positive cells were quantified per field from at least 3 sections per animal. Brain sections were collected between bregma −0.82 and −2.4 mm. Scale bars = 100 μm. Data are presented as mean ± SEM. *p < 0.05. Two-way ANOVA with Newman-Keuls post hoc test; p-values for the main effect of diet are indicated.

## DISCUSSION

Canagliflozin extends male but not female lifespan at 180 ppm, with male-specific neuroprotective effects, despite females accumulating higher drug concentrations in blood and brain (Miller *et al*. 2020; Jayarathne *et al*. 2022). Here we report that a subclinical concentration of Cana at 60 ppm improves glucose tolerance in both sexes but fails to reproduce the robust cognitive or neuroprotective profile observed at 180 ppm in either sex. Notably, the neurobiological response to low-dose treatment differed from that previously reported at the higher dose: while cognitive benefits were absent in both sexes, females exhibited reduced age-associated hippocampal microgliosis and astrogliosis, a phenotype not observed at 180 ppm, where neuroprotection was largely restricted to males. Our findings suggest that the central actions of Cana are both dose- and sex-dependent and further support the conclusion that differences in drug accumulation or concentration cannot explain sex-specific responses to Cana. Our results point to differences in CNS responsiveness as a key determinant of the sex-specific neurobiological outcomes observed between both sexes.

The sex-specific neuroprotective effects of Cana raise two possibilities. First, Cana may activate pathways particularly relevant to the male brain that are less responsive or differently regulated in the aged female brain (Healy *et al*. 2024; Zhang *et al*. 2025). This is consistent with the profiling of the aging hippocampus, which showed that 180 ppm Cana induces cyclic GMP-dependent signaling across transcriptomic, metabolomic, and proteomic layers in males but not in females (Jayarathne *et al*. 2025). Similarly, Acarbose, another ITP longevity intervention that preferentially extends lifespan in males, reduces neuroinflammation in males but not in females (Harrison *et al*. 2014; Sadagurski *et al*. 2017), suggesting that male-specific brain responsiveness to metabolic interventions applies to at least two agents, perhaps because both lower peak plasma glucose. Alternatively, Cana may produce equivalent primary benefits in both sexes but also engage negative effects in female tissues at higher exposures. This idea was recently supported by the shortening of female lifespan when 180 ppm is initiated late in life, with higher drug accumulation in aged females (Miller *et al*. 2024).

Feeding animals with 60 ppm Cana led to improvements in glucose tolerance in both sexes at 18 months without significant changes in body weight, suggesting that the glycemic and body weight effects of SGLT2i can be dissociated, consistent with glycemic efficacy across a broad dose range independent of weight loss (Pinto *et al*. 2022). The absence of changes in energy expenditure, in contrast to the significant increase at 180 ppm (Jayarathne *et al*. 2024), suggests that broader metabolic reprogramming may be needed for full neuroprotection. In support, rapamycin similarly shows dose-dependent metabolic reprogramming following its lifespan-extending effects (Miller *et al*. 2014). Whether the observed metabolic benefits at 60 ppm are sufficient to drive a lifespan effect in the absence of broader metabolic reprogramming remains to be seen in the ongoing ITP lifespan study https://www.nia.nih.gov/research/dab/interventions-testing-program-itp/supported-interventions.

The absence of cognitive improvements at 60 ppm in either sex supports a dose-threshold model of CNS neuroprotection. The reduced microgliosis in CA3 and DG without cognitive benefit in the female hippocampus is consistent with previous work showing that suppression of neuroinflammation alone is insufficient for cognitive improvement (Ma *et al*. 2025). Neuroinflammation is one of several interacting pathological processes in brain aging, together with mitochondrial dysfunction, synaptic loss, and impaired insulin signaling (Mani *et al*. 2025). One additional caveat is that we assessed cognition at an earlier time point (18 months) and neuroinflammation at 24 months; testing at a later age could provide additional insight. In males, improvements in Barnes maze performance and reductions in hippocampal gliosis were observed at 180 ppm but not at 60 ppm (Jayarathne *et al*. 2022; Jayarathne *et al*. 2025), suggesting that both effects are part of the same dose-dependent neuroprotective program. The lack of cognitive benefit in females despite partial reduction in gliosis may indicate that low-dose SGLT2i engages different brain targets in females, perhaps through pericyte-mediated mechanisms that operate independently of systemic glucose lowering (Takashima *et al*. 2022), but this remains to be further investigated.

The region- and cell-type-specific expression of SGLT2 in the brain may provide an additional explanation for the sex- and region-specific neuroinflammatory responses. Although SGLT2 was originally characterized as a renal transporter, evidence demonstrates its expression in neurons of the cortex, hippocampus, hypothalamus, and amygdala (Mei *et al*. 2024; Yeh *et al*. 2026) as well as in pericytes at the blood-brain barrier (Yu *et al*. 2010) and, critically, in astrocytes under conditions of metabolic stress, where SGLT2 upregulation drives cytotoxic edema through Na+-coupled glucose uptake in GFAP-positive cells, effects inhibited by Cana (Shim *et al*. 2023). Whether sex differences in astrocyte SGLT2 inducibility or downstream signaling contribute to the differential neuroinflammatory responses between males and females remains an open question.

Our findings have direct translational relevance for the repurposing of SGLT2 inhibitors in cognitive aging and neurodegeneration. SGLT2i have shown neuroprotective effects in rodent models of Alzheimer’s disease (AD), including reduction of amyloid burden, suppression of NLRP3 inflammasome activation, and protection of dopaminergic neurons (Kostrzewska *et al*. 2025; Mazzeo *et al*. 2026), and Mendelian randomization studies suggest that SGLT2i may enhance cognitive function in humans (Chen *et al*. 2025). However, sex-stratified analyses of these outcomes are limited. A meta-analysis of eleven randomized controlled trials found that the relative risk reduction in primary cardiac outcomes with SGLT2i was significantly lower in women than in men (Shah *et al*. 2024). The sex-specific hepatic transcriptional responses, differential pharmacokinetics, and divergent CNS responsiveness identified in this and previous studies provide a framework for understanding the sexually dimorphic effects of Cana. Our findings emphasize the importance of both sex-specific analyses and dose optimization when considering SGLT2 inhibitors for repurposing as interventions for brain aging.

In summary, we demonstrate that the neuroprotective effects of Cana are both dose- and sex-dependent, revealing fundamental sex differences in the CNS and pharmacokinetic response to SGLT2 inhibition. The molecular basis for these sex-specific responses remains to be determined and will require paired transcriptomic, proteomic, and metabolomic profiling of brain tissue across doses and sexes. Whether these findings will ultimately translate into lifespan outcomes remains an important open question.

## Abbreviations

Cana: Canagliflozin
CNS: central nervous system
SGLT2i: Sodium-glucose co-transporter 2 inhibitors
EE: energy expenditure
RER: respiratory exchange ratio

## Acknowledgements

This study was supported by Impetus, RF1AG078170, R01ES033171 for M.S. B.C.G was supported by U01 AG022307 and P30 AG013319. S.S was supported by T32-ES036169. Services and support were provided by the WSU Microscopy, Imaging and Cytometry Resources Core P30CA22453 and R50CA251068-01 and WSU Genomics Core.

## Author contributions

H.S.M.J., D.H.M, S.S., N.D.H.M., L.D., O.K., and N.R. carried out the research and reviewed the manuscript. R.A.M. provided suggestions about data interpretation. B.C.G analyzed the Cana levels in the brain. M.S. designed the study and analyzed the data. M.S. wrote the manuscript and is responsible for the integrity of this work. All authors approved the final version of the manuscript.

## Data Availability Statement

Further information and requests for resources and reagents should be directed to and will be fulfilled by the Lead Contact, Marianna Sadagurski. Any additional information required to reanalyze the data reported in this paper is available from the lead contact upon request.

